# Will the Illinois chorus frog (*Pseudacris streckeri illinoensis*) survive in Arkansas? A case study of a secretive species In need of protection

**DOI:** 10.1101/338699

**Authors:** Malcolm McCallum, Stanley E. Trauth

**Affiliations:** School of Agriculture and Applied Sciences, Langston University, Langston, OK 73050

**Keywords:** Illinois chorus frog, *Pseudacris streckeri illinoensis*, Arkansas, range contraction, amphibian declines

## Abstract

The range of the Illinois chorus frog (*Pseudacris streckeri illinoensis*) in Arkansas is restricted to the eastern quarter of Clay County. Nearly 100% of this species’ native sand-prairie habitat has been converted to agricultural fields. The original range of the Illinois chorus frog encompassed at least 9,982 ha. Although two new localities were identified in 2002, the current range is only 4,399 ha in 2002. This represents a 56% range contraction since 1992. Calling was heard in only 44.5% of its original range. This species may be experiencing a severe range contraction. Decay models predict the extirpation of the Illinois chorus frog in Arkansas within 17.5 to 101 yr. Suggested factors contributing to this range contraction may include drought, pesticide use, changes in surface water hydrology, U.S. E.P.A. Best management practices, and this species’ limited ability to recolonize extirpated sites.

## Introduction

The more difficult an organism’s presence is to detect, the more likely that declines in that species can proceed without notice. This is an important problem because the ‘ecosystem services’ these organisms provide are an integral support system for our planets survival (Wilson, 1988). Many amphibians reside in recesses deep beneath talus slopes, gravel beds, or other fossorial habitats for at least part of the activity season, and they are already among the most extinction-prone groups on the planet (McCallum 2007; McCallum 2015). This component of their life histories makes them difficult to detect and their population status difficult to monitor. A case example of this problem is the Illinois chorus frog (*Pseudacris streckeri illinoensis*).

The Illinois chorus frog (*Pseudacris streckeri illinoensis*) leads a fossorial existence, being found on the surface during only its 3-4 week breeding season early February through mid-March. Monitoring efforts are complicated by the hesitancy of this species to call when nighttime air temperatures fall below 4.5° C. Its sensitivity to temperature can leave a calling window of only a few hours per night on a few selected nights within a breeding season (Trauth et al. 2004; McCallum et al. 2006). This requires an investigator to travel at short notice to search for choruses. Unless a researcher lives within driving distance to the choruses, population monitoring is nearly impossible. Access to breeding ponds is further complicated, because the entire metapopulation in Arkansas resides on private lands.

The Illinois chorus frog is endemic to disjunct sand prairies extending from Clay County in northeastern Arkansas, across the boot heel of southeastern Missouri, and northward along the Mississippi and Illinois rivers into Illinois (Conant and Collins, 1998; Trauth et al. 2004). The populations in Arkansas represent the most recent biogeographic connections between the Illinois chorus frog and Strecker’s chorus frog (*Pseudacris streckeri streckeri*), found in the Arkansas River Valley. The geographic separation of these taxa during the past 8,000 years (Smith, 1957) has generated questions as to the validity of designating these two frogs as the same species (Trauth et al. 2007). Consequently, some authors (e.g., Collins, 1997) have chosen to elevate the Illinois chorus frog to full specific status pending further study. The highly fragmented eastern metapopulations of both taxa are matters of concern and undoubtedly place them at high risk of extirpation.

The ecological requirements of the Illinois chorus frog are narrowly defined and, because of its secretive nature, its natural history is largely unknown. These factors place it at an elevated risk of extirpation. The Illinois chorus frog is confined to sand prairies possessing numerous vernal ponds for breeding. The early breeding season makes it susceptible to sudden freezes, and it may easily succumb to cold weather (Tucker, 2000). Sandy soils, combined with its unusual forward burrowing behavior (Brown, 1972), may facilitate its escape from freezing temperatures. Observances of frostbite scars (McCallum and Trauth, pers. observ.; Tucker, 2000) support the contention by Packard et al. (1998) that there still might be significant mortality after sudden freezes. The Illinois chorus frog is incapable of burrowing effectively through soils with high silt/clay components (Brown, 1972). The compacted nature of non-sandy soils can also cause injuries to these frogs as they burrow, thus providing a route for infection by pathogens (MLM, pers. observ.). Illinois chorus frog tadpoles are cannibalistic (McCallum and Trauth 2001). This combined with the highly ephemeral nature of their breeding ponds, suggests that annual breeding success may vary greatly. Cannibalism in tadpoles is recognized as an adaptation to an erratic environment where recruitment of offspring can be quiet variable (Crump, 1983, Polis, 1981). The more variability in recruitment, the more susceptible to extirpation a species will be (Pechmann, 2001). From this evidence we can deduce that the Illinois chorus frog may be highly susceptible to extirpation in Arkansas as well as elsewhere across its range.

Like spadefoot toads (*Scaphiophus* and *Pelobates*), hellbenders (*Cryptobranchus*), cave-dwelling salamanders (*Typholotriton* and *Troglodytes*), and other similarly vulnerable species, it is easy for their population trends to change without notice due to their extreme secretive nature or the accompanying inaccessibility of their habitats (Wilson, 1988). Frequent monitoring of these groups is necessary to safeguard against permanent loss of these unique taxa.

## Materials and Methods

We conducted a status survey of the Illinois chorus frog in 2002 primarily because of concerns about its well being in the state. Previously, Trauth conducted less intensive status surveys in 1985, 1986 and 1992. Informal visits to key ponds occurred every year between 1986 and 1992, and periodically between 1992 and 2002. In all but the 2002 survey, voucher specimens were collected for deposition into the Arkansas State University Museum of Zoology (ASUMZ) herpetological collection.

### 1985 Survey

This was the first attempt to accurately define the range of Illinois chorus frogs in Arkansas. This survey was conducted in areas containing sandy soils. Thirty potential breeding sites in five northeast Arkansas counties were surveyed on eight different dates from 22 February through 14 April 1985. All areas were visited at night just prior to, during, or after rainfalls on nights when air temperatures exceeded 10° C. Presence or absence and population estimates of male Illinois chorus frogs were recorded along with observations of sympatric species.

### 1986 and 1992 Surveys

Four counties in northeast Arkansas were searched on 11 different occasions from 18 January through 15 March 1986. Efforts were restricted to Clay County, Arkansas during surveys conducted between 11 February and 25 March 1992. Many areas were repeatedly visited on the same and different nights. All visits to potential breeding habitats were conducted during or just following rainy weather. The primary survey tool used was calling surveys. Surveys were conducted by stopping randomly along highways and farm roads and then listening for choruses. If calling male Illinois chorus frogs were heard, the location of the chorus was plotted on the map by hand. Additional observations of sympatric amphibian species were recorded.

### 2000 – 2002 Surveys

Breeding chorus surveys were conducted in Clay County, Arkansas on the following dates: February 18, 28 and March 4, 2000; February 13, 15, 24, 2001; February 14, 19, 23 and March 1, 8, 15, 2002. We followed sampling protocols deployed by the U.S. Geological Survey’s Biological Resources Division, North American Amphibian Monitoring Program [NAAMP]. Environmental parameters and presence of calling frogs were recorded during each visit. Surveys were conducted by road cruising through the eastern quarter of Clay County (Fig. 1) beginning at 2200 hr. County roads bordering at least two sides of each square cartographic Section were driven. Listening stations were plotted at the corner and midpoint of each side for each Section within a Township. Stations were inventoried by turning off the car, leaving the vehicle, and listening for a period of 3-5 minutes. If calling was heard at a location, it was recorded on a map. Daytime visits were conducted to determine the extent of breeding pools and assess condition of the habitat. Habitat condition was classified as follows:

1. Highly disturbed (i.e., no native habitat available, 100% agricultural fields).
2. Moderately disturbed (i.e., some prairie-like habitat evident, within visual vicinity).
3. Somewhat disturbed (i.e., prairie-like habitat is evident, either bordering or partially in farmland).
4. Undisturbed (i.e., prairie-like habitat, farmland is not within 1,000 m of the chorus).

**Figure 1.**
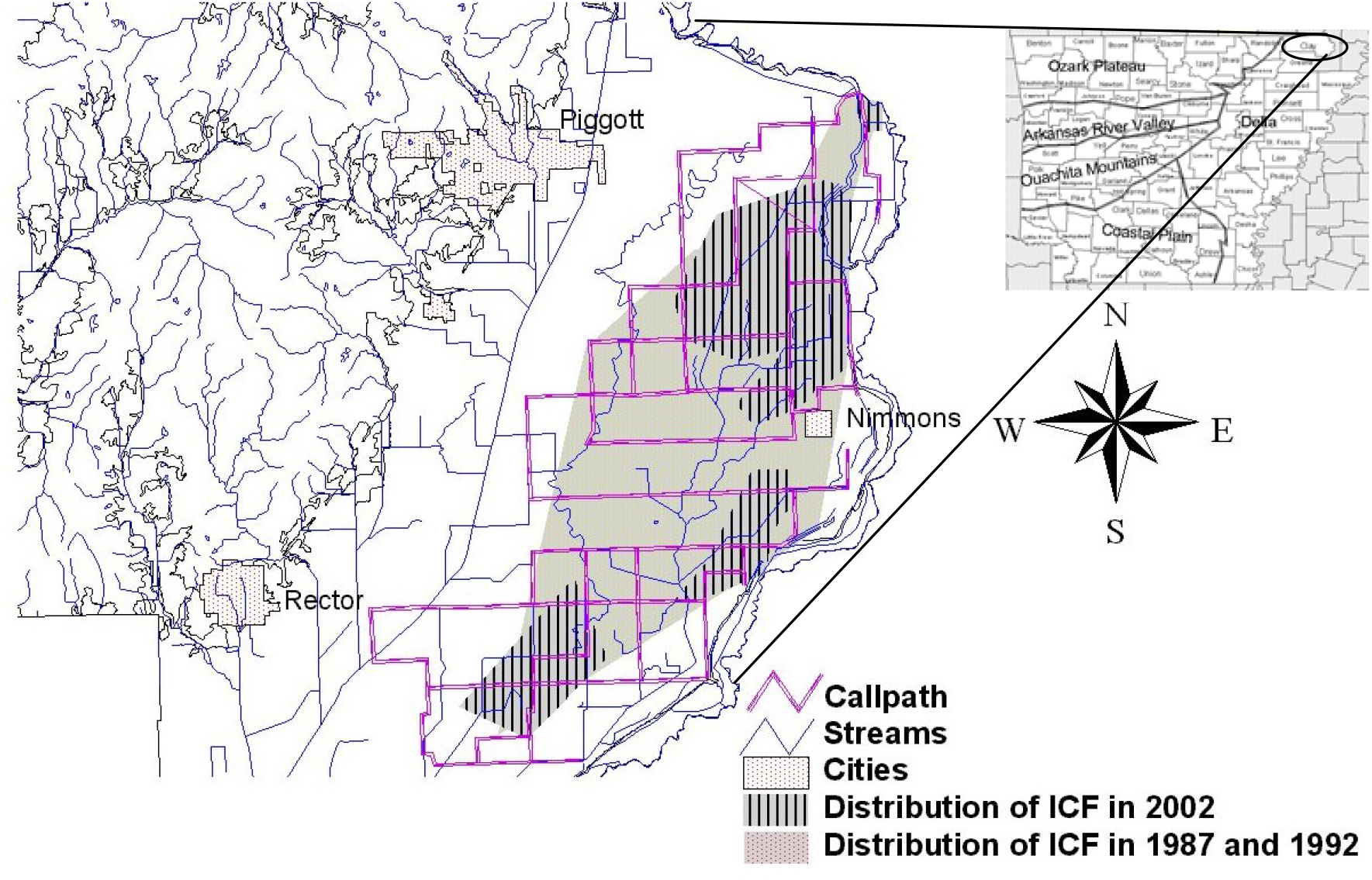
The Current and historic distribution of the Illinois chorus frog in Clay County, Arkansas.

The distribution of choruses was mapped using ArcView GIS 3.2. Range contraction was mathematically modeled using both linear decay (y = mx + b) and uninhibited logistic decay (d/Q = *k*Q) formulas to provide estimates of the imminent risk of extirpation (Ricciardi and Rasmussen, 1999).

## Results

### 1985 Survey

The Illinois chorus frog was found at only three localities, all in Clay County. Male specimens (ASUMZ 4177 and 4178) were collected and deposited in the museum. All three localities were confined to Middle Slough, a tributary of the St. Francis River. No choruses were identified in the Western Lowlands. This region is the portion of the Mississippi Embayment located west of Crowley’s Ridge and East of the Ozark Plateau (Saucier, 1978). Calling was observed in Clay County, Arkansas on three dates (Table 1).

**Table 1.** Calling dates for Illinois Chorus Frogs in Clay County, AR from 1986-2002.

| Dates | Calling (yes/no) |
| --- | --- |
| 22 February 1985 | Yes |
| 22 March 1985 | Yes |
| 29 March 1985 | Yes |
| 18 January 1986 | No |
| 2 February 1986 | No |
| 18 February 1986 | Yes |
| 8 March 1986 | No |
| 12 March 1986 | Yes |
| 14 March 1986 | Yes |
| 15 March 1986 | Yes |
| 18 February 2000 | No |
| 28 February 2000 | Yes |
| 4 March 2000 | Yes |
| 13 February 2001 | No |
| 15 February 2001 | Yes |
| 24 February 2001 | Yes |
| 14 February 2002 | No |
| 19 February 2002 | Yes |
| 23 February 2002 | Yes |
| 1 March 2002 | No |
| 8 March 2002 | Yes |
| 15 March 2002 | Yes |

### 1986 Survey

A large breeding chorus of more than 50 males was observed in a pond ca. 125 m^2^ located ca. 2.4 km northwest of Nimmons (Clay County, Arkansas). Breeding spadefoot toads (*Scaphiopus holbrookii*) were also observed at this site. The 1986 distribution of Illinois chorus frogs was expanded to encompass 2,130 ha (Fig. 1). The three calling locations identified in 1985 did not possess calling in 1986. A significant amount of habitat alteration had occurred through changes in cultivation practices at those sites. Calling was observed from 18 February through 15 March (Table 1).

### 1992 Survey

The known range of the Illinois chorus frog was expanded by 78% to 9,892 ha (Fig. 1). Nearly all breeding populations were located in or along cultivated fields. All localities were associated with Middle and Hampton sloughs. Grass line habitats along fencerows and slough drainages showed little resemblance to the natural prairie habitat but represent the majority of “non-cultivated” habitat available. A single large breeding site near Nimmons has been used every year from 1986 to 1992. This location appeared to have entered cultivation in 1991. This site also contained at least 20 males in 1992. All sites with calling in 1986 possessed calling in 1992.

### 2000-2002 Surveys

#### Abnormalities

A single male with a missing arm was observed during the entire spring 2000 chorus. Apparent frostbite scars (Tucker, 2000) were observed in 10 individuals during 2001. Two more possessed red inguinal pustules (possibly from frostbite), one had localized lymphadema, and another individual had a dysfunctional vocal sac. No abnormalities were observed in 2002.

#### Environmental Variables and Calling

The pond located near Nimmons contained 10-15 calling males in 2000. In 2001 and 2002 no calling occurred at that site. On February 15, 2001 we observed 30 males and 10 females at a second primary breeding site.

Calling males were heard from February 28 through March 15, 2002 (Table 1). Breeding chorus size was not determined due to difficulties arising from varying distances between the observer and the breeding choruses. All choruses fell within NAAMP category 2, calls of individuals can be distinguished with overlapping calls. Climatic conditions, when choruses were not heard, were characterized by ambient air temperatures < 14° C and wind speeds greater than 4.8 km/hr. Climatic conditions when choruses were heard were characterized by ambient air temperatures > 14° C and wind speeds less than 1.6 km/hr. On only one occasion (March 8, 2002) were males observed calling below 14° C.

#### Range Contraction

In 2002 range encompassed only 4,400 ha in Clay County, with two previously unknown chorus sites identified (Fig. 1). This represents nearly a 56% range contraction since 1992. Only 57% of the range (1,222 ha) identified in 1987 possessed choruses in 1992. Assuming all 11,068 ha known to have contained choruses did so in 1987, it appears that the Illinois chorus frog has experienced a range contraction, as only 39% of its previously mapped range contained frogs in 2002 (Fig. 2). All active choruses were found in highly disturbed sites, habitat classification Code 1 (Fig. 3).

**Figure 2.**
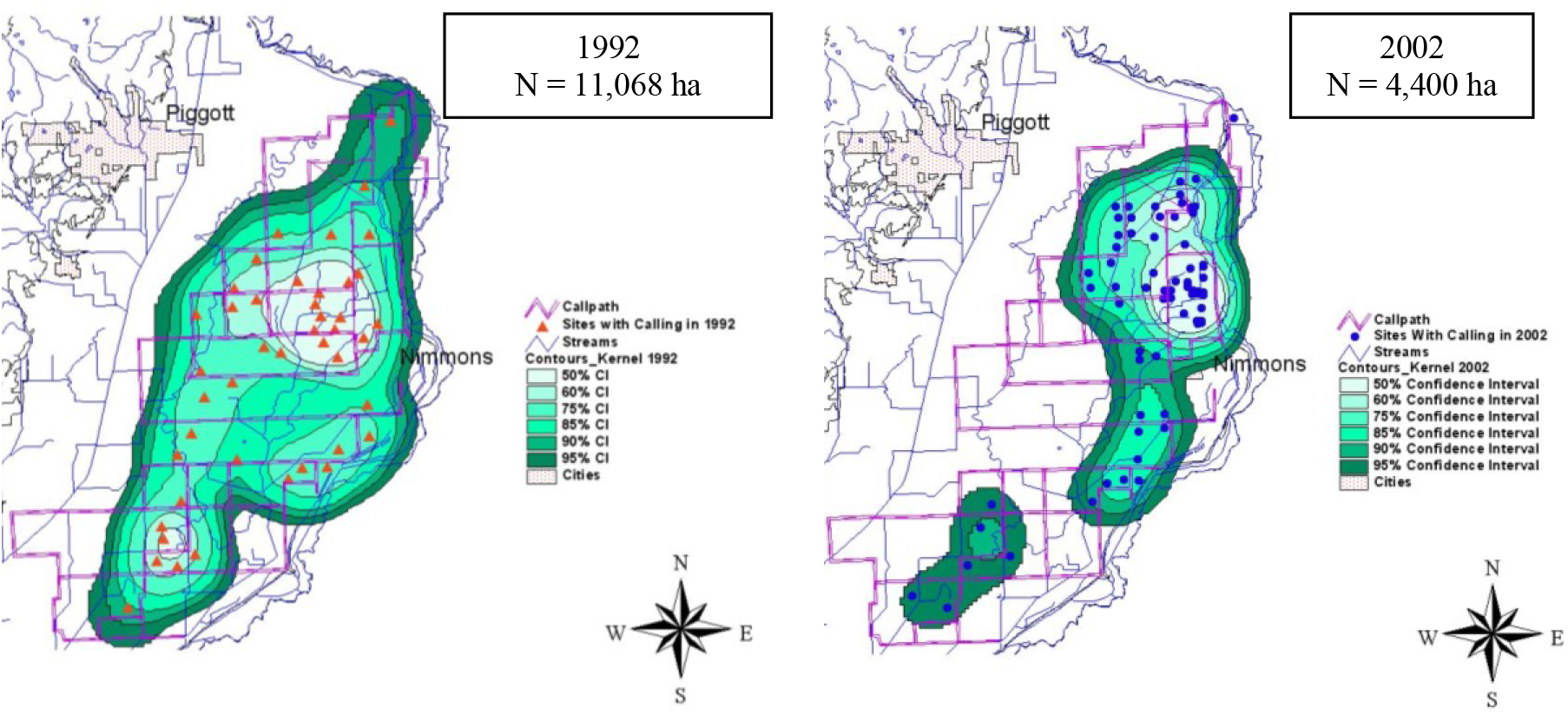
Range contraction of the Illinois Chorus Frog (*Pseudacris streckeri illinoensis*) in northeastern Arkansas from 1992 – 2002.

**Figure 3.**
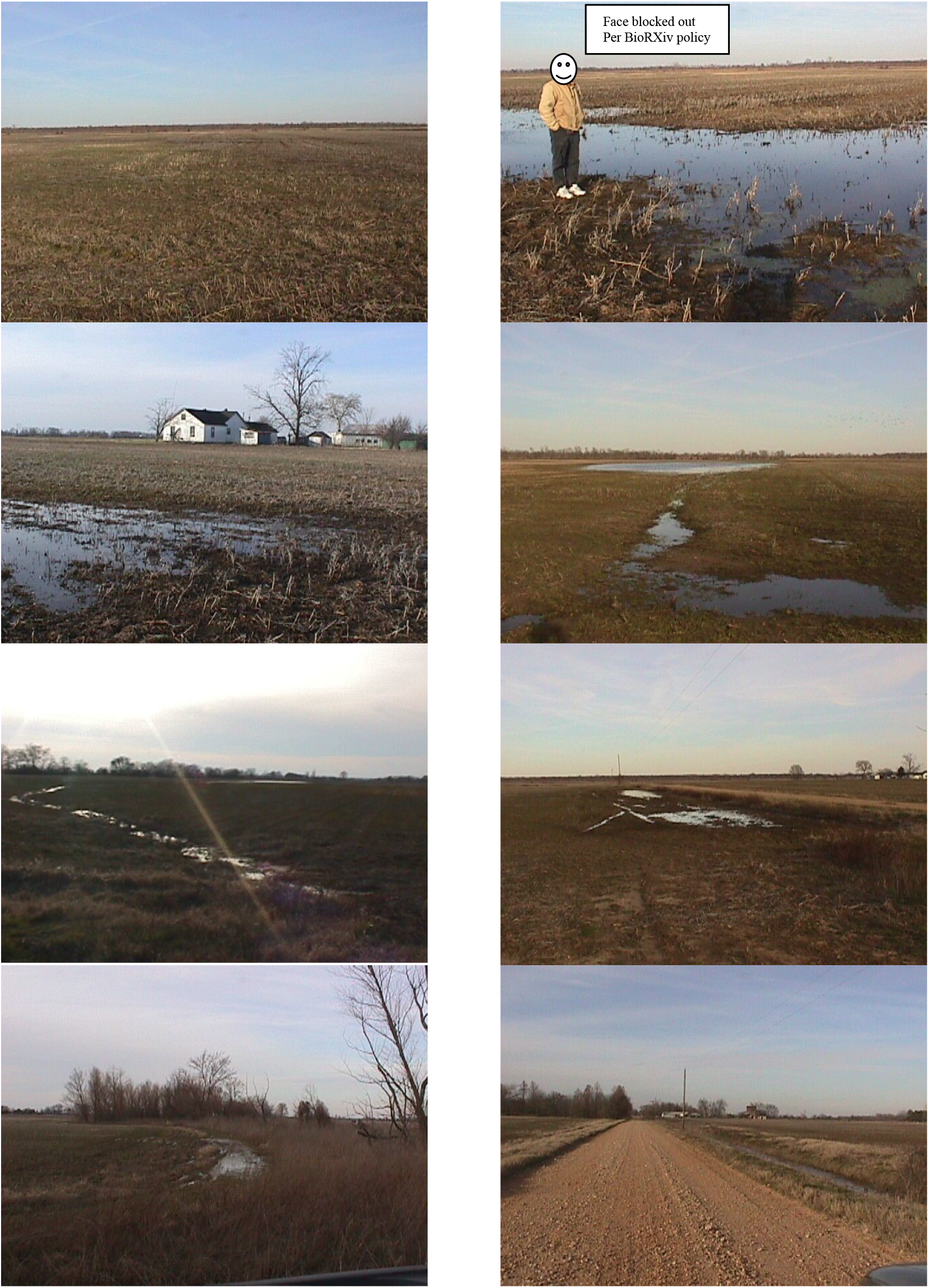
Photographs of breeding habitat condition in Clay County, Arkansas in 2002.

#### Modeling

Using the maximum number of hectares (n = 11,068) containing choruses from 1987 – 2002, as the original distribution in Clay County, the linear decay model suggests the Illinois chorus frog will be extirpated from Arkansas within 17 yr. The uninhibited logistic decay model predicts the range will continue to contract slowly and persist for 101 yr.

## Discussion

Cultivated fields dominate the landscape of eastern Clay County, and few filter strips, field borders, or hedgerows are present. In addition, drainage ditches to prevent pooling of water bound most fields. The entire Arkansas population resides within a landscape predominated by cotton and soybean fields. Tucker (1997) found cutworms, a common agricultural pest, to be a primary food of this species. Because of the intensive application of pesticides to control cutworms and other agronomic pests, this anuran is probably routinely exposed to pesticides.

The brief breeding season of the Illinois chorus frog presents several problems for future monitoring efforts. When attempting to monitor or conduct research on Illinois chorus frog choruses in Arkansas, visitation to breeding sites should begin after the first week of February, depending on weather conditions. If there is unseasonably warm weather earlier than this, the frogs may begin breeding in January. Based on our observations chorusing normally occurs sporadically for about four weeks. Wind can be a significant factor when monitoring Illinois chorus frog. Chorusing appears to be suppressed by windy conditions. The high-pitched call is very weak and does not carry for long distances and calling is easily masked by strong or gusty winds. Because of this, it might be more informative to utilize the chorus characterization methods recommended by Lips et al. (2001) than NAAMP protocols when monitoring Illinois chorus frogs. These observations suggest that calling surveys are less effective under windy conditions. Similar problems were observed during rainy nights, particularly if the rain varied in intensity. Males ceased calling whenever rainfall intensity was significantly punctuated.

Temperature appears to greatly influence chorusing activity. Choruses were generally observed on nights when the ambient air temperature was > 14° C. The one exception to this occurred on a night following unseasonably warm daytime temperatures, and the nighttime temperature fell below 14° C. Based on this observation daytime temperatures can influence calling activity independently of nighttime temperatures by heating of the breeding pond water.

The apparent loss of previously known populations and current range contraction raises concerns about the viability of current populations and their potential ability to recolonize previously occupied sites. A minimum viable population (MVP) for any given species is the smallest isolated population having a 99% chance of remaining extant for 1,000 yr despite foreseeable demographic, environmental, or genetic effects (Primack 1998, Shaffer 1981). Our information suggests that the Illinois chorus frog may be extirpated in Arkansas in less than 100 years. This indicates that it is declining at a much more rapid rate than has been estimated for most other North American amphibians (Ricciardi and Rasmussen, 1999). It is also apparent that our current lack of conservation efforts to protect this species must be rectified.

Previous research suggests that small mammal reserves require at least 100 km^2^ for minimum viable area (MVA; Schonewald-Cox et al., 1983). The entire Arkansas Illinois chorus frog metapopulation currently occupies 38 km^2^ and has never been known to be larger than 59 km^2^. We have recommended that at least 6.4 km^2^ be placed in such conservation programs, this would represent almost one third of the remaining Illinois chorus frog range in Arkansas.

The range contraction we observed in this species went completely unnoticed despite frequent visits to the vicinity from 1992 through 2001. This suggests that species like the Illinois chorus frog are difficult to detect during most of the year; this also represents a serious problem if non-intensively monitored. This is especially true for populations that persist on private lands. Species that are difficult to detect and monitor are at an extreme risk of extirpation when they occur on private lands. Little monitoring of herpetofauna and other non-game taxa normally takes place on private lands. The monitoring that does take place is typically minor because of a significant lack of funds for this type of research. Funneling money into these kinds of activities by federal and state governments so that conservation efforts can become more proactive and less reactionary seems a worthwhile endeavor.

The change in Illinois chorus frog populations undoubtedly resulted from multiple contributing factors. First, Illinois chorus frogs presumably resist migrating between ponds reducing the potential for recolonization. Second, surface water hydrology might have changed in the last 10 yr as a result of aquifer draw down changing the periodicity and location of potential breeding pools. Finally, changes in land use and application of precision leveling via U.S. E.P.A. Best management practices for erosion control were implemented 1992 (Trauth et al. 2006). Remarkably, precision leveling accelerated after a public meeting with stakeholders in the region. Considering this species’ habitat requirements, possibly variable recruitment, probable exposure to contaminants, and expression of immuno-physiological disorders (McCallum 2018), it is essential that more studies be implemented to investigate options to preserve the Illinois chorus frog in Arkansas and elsewhere throughout its range.

Other species in this region, such as spadefoot toads, have not been observed in many years. Might they have been extirpated in this region? The current status of the Illinois chorus frog in Arkansas does provide an important notion to ponder. Just how many common but seldom seen species are disappearing in the backyards of North America? Hopefully, lands will become available to save the Illinois chorus frog in Arkansas, but what are we to do about those species currently disappearing that are not monitored?

## Acknowledgements

We thank B. Wheeler, B. Neal, C. McDowell, J. Varner, J. Jackson, B. Samples, and J. Gore for field assistance, Arkansas State University for the use of equipment and facilities, and the Arkansas Game and Fish for permits. This research was funded by the United States Geological Survey and the Arkansas Game and Fish.

